# Preclinical Characterization of Hippo pathway inhibition: Insights from Pharmacological and Genetic Studies

**DOI:** 10.64898/2026.08.01.742182

**Authors:** Sayantanee Paul, Michelle Lepherd, Thijs Hagenbeek, Satoko Kakiuchi-Kiyota, Mioaran Ning, Minyi Shi, Bence Daniel, Ryan Ybarra, Jessica Sims, Anwesha Dey

**Affiliations:** Department of Discovery Oncology; Translational Safety; Drug Metabolism and Pharmacokinetics; Proteomic and Genomic Technologies, Genentech Inc, South San Francisco, California, CA

## Abstract

The Hippo pathway is an evolutionarily conserved regulator of growth, regeneration, and organ homeostasis, and while its dysregulation is well established in cancer, the effects of inhibiting this pathway on normal tissues are less understood. Here we have systematically investigated the impact of Hippo pathway inhibition by comparing pharmacologic perturbation using a covalent small-molecule TEAD inhibitor (TEADi CMPD1, also known as GNE-8025) with genetic suppression of YAP/TAZ. We identified three key target organs that consistently emerged upon TEAD inhibition: the kidney, as well as the pancreas, and thymus. Across models, both perturbations led to comparable disease phenotypes in these organs, including tubular degeneration in the kidney, acinar atrophy in the pancreas, and lymphoid depletion in the thymus. However, the extent of damage was more pronounced in mice treated with the small-molecule inhibitor, highlighting potential dose and compound specific effects while remaining broadly consistent with the phenotypes observed upon genetic ablation of YAP/TAZ. This highlights the key role of evaluating both genetic and pharmacological perturbations to characterize the phenotypes and potential toxicities when modulating novel targets in oncology. To further investigate the mechanisms underlying pan-TEAD inhibition and kidney related adverse effects, we further characterized this class effect through a comprehensive transcriptomic analysis of the kidney to map the pathways involved in renal response.

**Significance:** Understanding on target toxicities is critical for the safe clinical development of TEAD inhibitors. Here, by integrating pharmacologic TEAD inhibition with genetic suppression by developing a mouse model that characterizes systemic, inducible knockdown of YAP/TAZ, we provide a systematic framework to define the Hippo pathway liabilities *in vivo*. We identify kidney, pancreas, and thymus as conserved target organs with concomitant phenotypes across both genetic and pharmacological methods, establishing these as pathway driven effects. Importantly, we uncover dose dependent and partially irreversible injury, particularly in kidney and pancreas, alongside mechanistic insight linking TEAD inhibition to aldosterone signaling disruption in kidney. These findings highlight the importance of strategies to identify monitorable, manageable adverse effect to guide clinical translation of TEAD targeting strategies.

## Introduction

The Hippo pathway is an ubiquitously expressed, evolutionarily conserved signaling cascade that regulates cell proliferation, tissue growth, organ development, and regeneration by suppressing excessive growth (1, 2). Central to this pathway is a kinase cascade involving MST1/2 (Mammalian Ste20-like kinases 1 and 2) along with their adaptor proteins SAV1, LATS1/2, and MOB1 (3, 4). TEAD transcription factors (TEAD1-4) play key roles in gene regulation but require YAP (Yes-associated protein) and TAZ (transcriptional coactivator with a PDZ-binding motif) as coactivators (5, 6), as TEADs lack intrinsic transcriptional activity (7). When the Hippo pathway is inactive, YAP and TAZ translocate to the nucleus, bind TEADs, and drive gene expression programs that promote cell proliferation and survival (8, 9). In contrast, activation of the Hippo pathway occurs through upstream tumor suppressors such as NF2/Merlin and KIBRA, which activate MST1/2, leading to the phosphorylation of LATS1/2 (1). Active LATS1/2 then phosphorylates YAP and TAZ, retaining them in the cytoplasm to be subsequently degraded, thereby preventing their nuclear activity (10–12). This regulatory mechanism is crucial for maintaining tissue homeostasis and is frequently disrupted in diseases such as cancer (13). Aberrant Hippo signaling has been implicated in multiple cancer types, including lung, liver, breast, skin, and colon cancers, where it is associated with poor prognosis and therapy resistance (13–15). Mutations in upstream components such as NF2 and LATS1/2, as well as YAP/TAZ overexpression, amplification, and nuclear localization, contribute to tumor initiation, progression, and metastasis (15–17). Emerging evidence suggests that Hippo pathway dysregulation cooperates with oncogenic signaling pathways such as KRAS, MAPK, and WNT to promote tumorigenesis and therapy resistance by transcriptional rewiring (17–23). The sustained activation of YAP and TAZ in both untreated and therapy-resistant cancers underscore the Hippo pathway as a promising therapeutic target (24, 25), driving academic and industry efforts to develop TEAD inhibitors (26) (27–30).

However, targeting the Hippo pathway presents challenges, particularly when inhibiting the TEAD transcription factors. Given the role of YAP/TAZ/TEAD in development, regeneration, and homeostasis, disrupting their function could lead to adverse effects in normal tissues. Insights from human genetic variants, rodent knockout models, and preclinical studies indicate that systemic TEAD inhibition may result in potential toxicities, and kidney specific adverse effects are emerging as TEAD inhibitors are progressing to the clinic (31–37). Preclinical and clinical data suggest that kidney-related adverse effects could be a common characteristic of pan-TEAD inhibitors the impact of which is further compounded by the limited understanding of the distinct and overlapping roles of TEAD paralogs in organ development and function (31, 33–38).

Here we show the effects of YAP/TAZ perturbation, through either genetic knockdown of YAP/TAZ or by targeting TEAD with a small molecule inhibitor, on major organs like the kidney, pancreas, and thymus. TEAD proteins contain a cysteine residue (Cys380) within the lipid pocket that is normally modified by endogenous palmitoylation (26, 27). This cysteine is conserved across all four TEAD paralogs and can be covalently engaged by small molecules to allosterically disrupt the interaction between TEAD and YAP (29, 39). In this context, we used TEADi CMPD1 (also known GNE-8025), a potent covalent inhibitor that strongly engages the TEAD lipid pocket across all isoforms except TEAD3 where there is a more modest engagement compared to TEADs 1, 2, and 4. TEADi CMPD1 demonstrates robust pathway inhibition and anti-tumor efficacy in Hippo dysregulated models (40). In summary we provide characterization of the pharmacological and genetic perturbation of YAP/TAZ/TEAD dependent responses of the Hippo pathway. Targeting this pathway shows substantial kidney injury as well as effects in other organs including pancreas and thymus. Preclinical and clinical studies suggest that some adverse effects are potentially monitorable, manageable, and reversible in the clinic, continued vigilance and careful safety monitoring will remain essential as these agents advance through clinical development.

## Results

### Pharmacological and genetic inhibition of YAP/TAZ *in vivo*

To investigate how systemic inhibition of YAP/TAZ signaling impacts tissue homeostasis, we employed both pharmacological and genetic perturbation strategies in mouse models. For pharmacologic inhibition, Crl: CD1 mice were treated with TEADi CMPD1, a covalent palmitoylation-site binder, for 28 days at doses of 0, 10, 50, or 250 mg/kg/day (Figure 1A) with a cohort of animals from the 0, 50 and 250 mg/kg groups followed for another 28 days after dosing cessation to assess recovery of any effects. Dose dependent increase in systemic exposure to the TEADi was confirmed on Day 1 and Day 28 of the study (Figure S1A). In parallel, we used a doxycycline-inducible shRNA system to systemically deplete YAP and TAZ, with doxycycline administered for 14 days before analysis. We confirmed the knockdown in this study by assessing YAP/TAZ protein levels via immunohistochemistry (Figure S1B, C), which was consistent with our prior work demonstrating pathway modulation upon YAP/TAZ knockdown in the same mouse model (41). Across both pharmacologic and genetic models, we evaluated a comprehensive panel of endpoints, including serum chemistry, hematology, urinalysis, organ weight ratios, and histopathology of major tissues. These approaches enabled us to characterize how the pathway inhibition affects tissue architecture and organ function *in vivo*.

**Figure 1.**
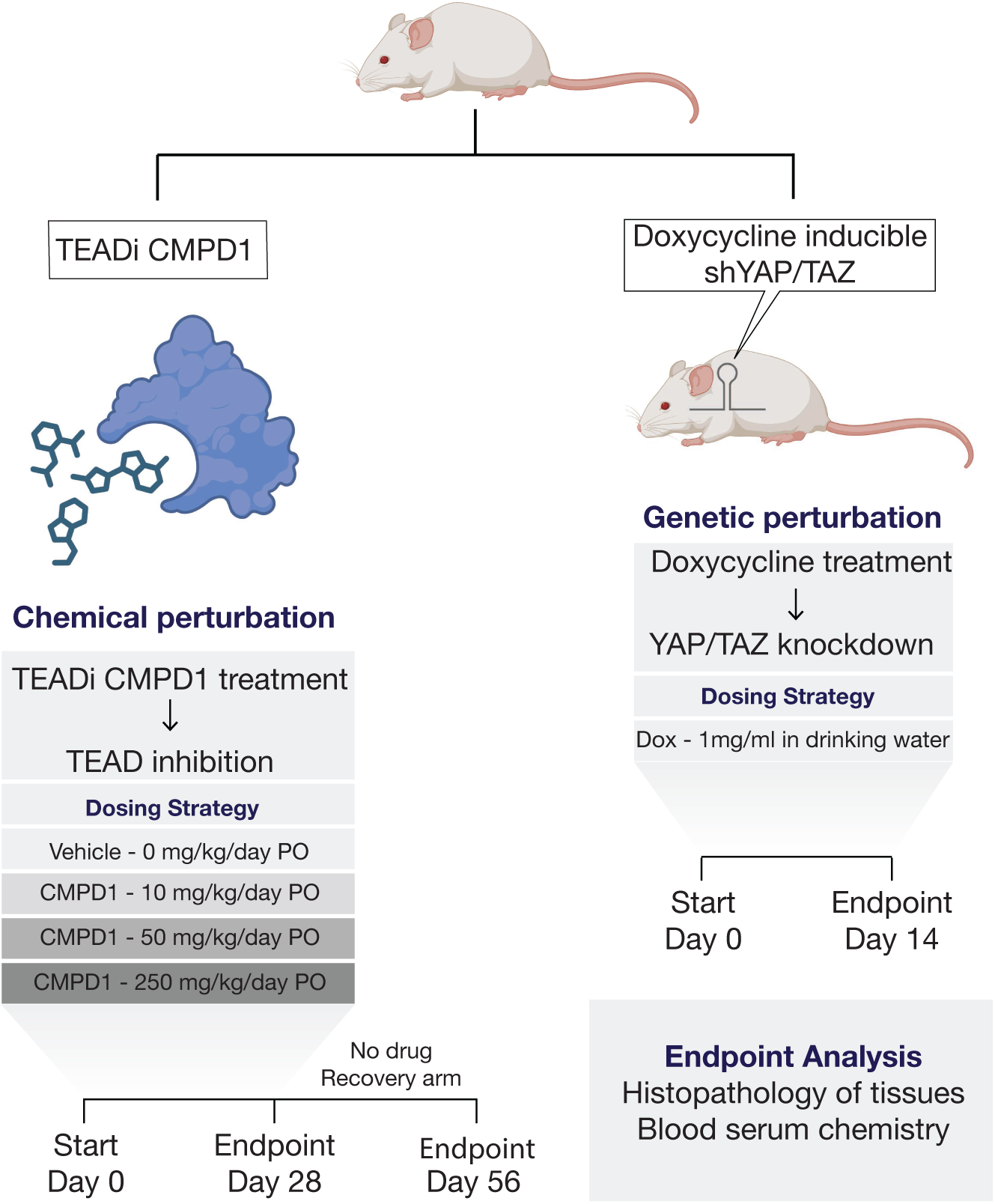
Experimental strategy for pharmacological and genetic inhibition of YAP/TAZ *in vivo*. Schematic of two experimental approaches used to inhibit YAP/TAZ activity in vivo. Left: For chemical perturbation, mice were treated with the TEAD palmitoylation inhibitor TEADi CMPD1 at doses of 0,10, 50, 250 mg/kg, once daily for 28 days by oral gavage (PO). In a separate cohort, animals were followed for an additional 28-day drug-free recovery period after completion of TEADi CMPD1 dosing. Right: Mice expressing a doxycycline-inducible shRNA targeting both YAP and TAZ as well as wildtype controls were given 1 mg/ml doxycycline in 5% sucrose water for 14 days to induce genetic knockdown. For both experiments, blood samples and tissues were collected at the end of the treatment period for clinical pathology and histological analysis.

### YAP/TAZ inhibition induces dose-dependent renal findings

To assess the impact of YAP/TAZ inhibition on the kidney, we performed histopathological and clinical pathology analyses following both pharmacological and conditional genetic perturbations. Administration of TEADi CMPD1 for up to 28 days in Crl:CD1 mice resulted in dose-dependent adverse effects in the kidney primarily affecting the tubules (Figure 2A). The highest dose tested (250 mg/kg) was not tolerated and resulted in death or the need for early euthanasia of the animals between days 17 and 28 of dosing. The primary clinical observations included decreased mean body weight (14.1% and 10.4% lower than control for males and females, respectively), decreased food consumption (15.2% and 18.0% lower than control for males and females, respectively), decreased activity, recumbency, bradypnea and/or hypothermia. Prior to euthanasia, a severe increase in urine volume was noted at doses of ≥ 10 mg/kg and correlated with histologic findings, primarily in the corticomedullary region of the kidney, of tubular dilatation and degeneration, hyaline casts/proteinosis, fibroplasia, and tubular regeneration. At 250 mg/kg, there was also a correlating increase in the kidney to brain weight ratio (Figure 2B). Additional clinical pathology findings associated with adverse effects related to the kidney included decreased urine specific gravity and increased serum inorganic phosphate (data not shown), sodium and chloride at ≥ 10 mg/kg; increased serum creatinine and urea, decreased serum albumin, and increased urine protein: creatinine ratio at ≥ 50 mg/kg; and increased serum potassium at 250 mg/kg (Figure 2B). In the transgenic mice with 14 days of doxycycline-induced dual knockdown of YAP and TAZ similar, although milder, tubular dilatation, degeneration, and hyaline casts were observed microscopically in the kidney (Figure 2C) and the more limited clinical pathology profile tested showed similar trends with increases in urea, creatinine, sodium, chloride, and potassium and decreases in albumin (Figure S1D), providing additional supportive evidence that the renal adverse effects observed in the TEADi CMPD1 treated mice was a direct consequence of YAP/TAZ suppression. Together, these results demonstrate that systemic inhibition of YAP/TAZ signaling leads to adverse effects in kidney in a dose-dependent manner.

**Figure 2.**
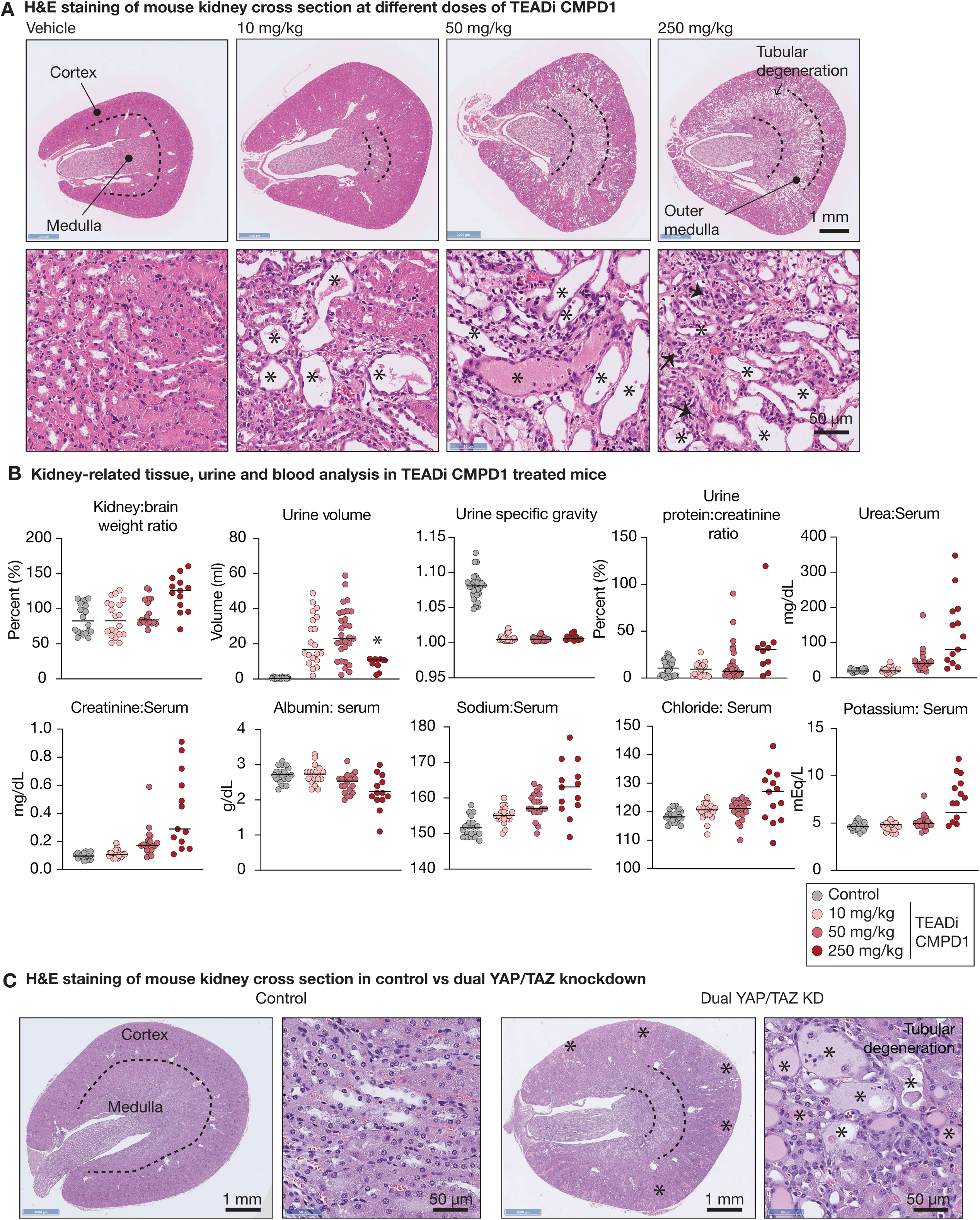
YAP/TAZ inhibition induces renal tubular degeneration. (A) H&E staining of mouse kidney showing tubular dilatation and degeneration, primarily in the outer medulla zone of the kidney after 28 days of vehicle or TEADi CMPD1at 10 mg/kg and 50 mg/kg and after 22 days of TEADi CMPD1 at 250 mg/kg. Dotted line on low magnification vehicle image is approximate location of corticomedullary junction; Dashed lines at ≥ 10 mg/kg border the approximate zones of the most severe tubular degeneration in the outer medulla; Asterisks = dilated tubules with attenuated or sloughed epithelial cells +/- intratubular cellular and proteinaceous debris; arrow heads = areas of tubular loss and interstitial replacement by early fibroplasia, most prominent at 250 mg/kg. Scale bars: 1 mm (top panels), 50 µm (bottom panels). (B) Kidney weight, urine analysis, and clinical chemistry in mice after 28 days of vehicle or TEADi CMPD1 at 10 mg/kg and 50 mg/kg and after 17 to 24 days of TEADi CMPD1 at 250 mg/kg. There is increase in kidney weight (measured as the kidney to brain weight ratio to account for body weight loss) in the 250 mg/kg dose group. Urine volume is severely increased and urine concentration (as measured by urine specific gravity) is markedly decreased in all treated animals (≥ 10 mg/kg); note for animals in the 250 mg/kg group marked with an asterisk, the increase in urine volume was unexpected and the maximum volume in the collection pan was 11mL (for the 10 and 50 mg/kg dose groups, a larger collection pan was used to get a more accurate measurement). Urine protein (as measured by urine protein:creatinine ratio), serum urea, and serum creatinine are increased in the 50 mg/kg and 250 mg/kg dose groups. Serum albumin is decreased in the 250 mg/kg group and, to a lesser extent, the 50 mg/kg dose group, correlating with the increase in urine protein:creatinine ratio. Serum sodium and chloride are increased in the ≥ 10 mg/kg dose groups and potassium is increased in the 250 mg/kg dose group. Dots represent individual animals; horizontal lines indicate the median. (C) H&E staining of kidneys from wildtype control and dual YAP/TAZ knockdown mice showing tubular dilatation and degeneration in the cortex and outer medulla zone of the kidney after 14 days of doxycycline administration, confirming that renal damage results from loss of YAP/TAZ activity. Dotted line on wildtype image is approximate location of corticomedullary junction; Dashed lines in Dual YAP/TAZ KD kidney border the approximate zones of the most severe tubular degeneration in the outer medulla; Asterisks = dilated tubules with attenuated or sloughed epithelial cells +/- intratubular cellular and proteinaceous debris. Scale bars: 1 mm (left panels), 50 µm (right panels).

### YAP/TAZ suppression causes acinar cell injury and atrophy in the exocrine pancreas

Administration of TEADi CMPD1 for up to 28 days in Crl:CD1 mice resulted in dose-dependent pancreatic findings affecting the acinar cells of the exocrine pancreas (Figure 3A). At ≥ 10 mg/kg, there was multifocal vacuolation, degeneration, and atrophy of the acinar cells which was often associated with severe increases in serum amylase and lipase (Figure 3A, B). The endocrine pancreas (islets) were microscopically unaffected. A similar, although less severe, phenotype was observed in the exocrine pancreas of transgenic mice with 14 days of doxycycline-induced dual knockdown of YAP and TAZ where there was multifocal loss of zymogen granules and mild atrophy of acinar cells (Figure 3C). These findings demonstrate that both pharmacological and conditional genetic inhibition of YAP/TAZ disrupt pancreatic homeostasis, particularly of the exocrine cells.

**Figure 3.**
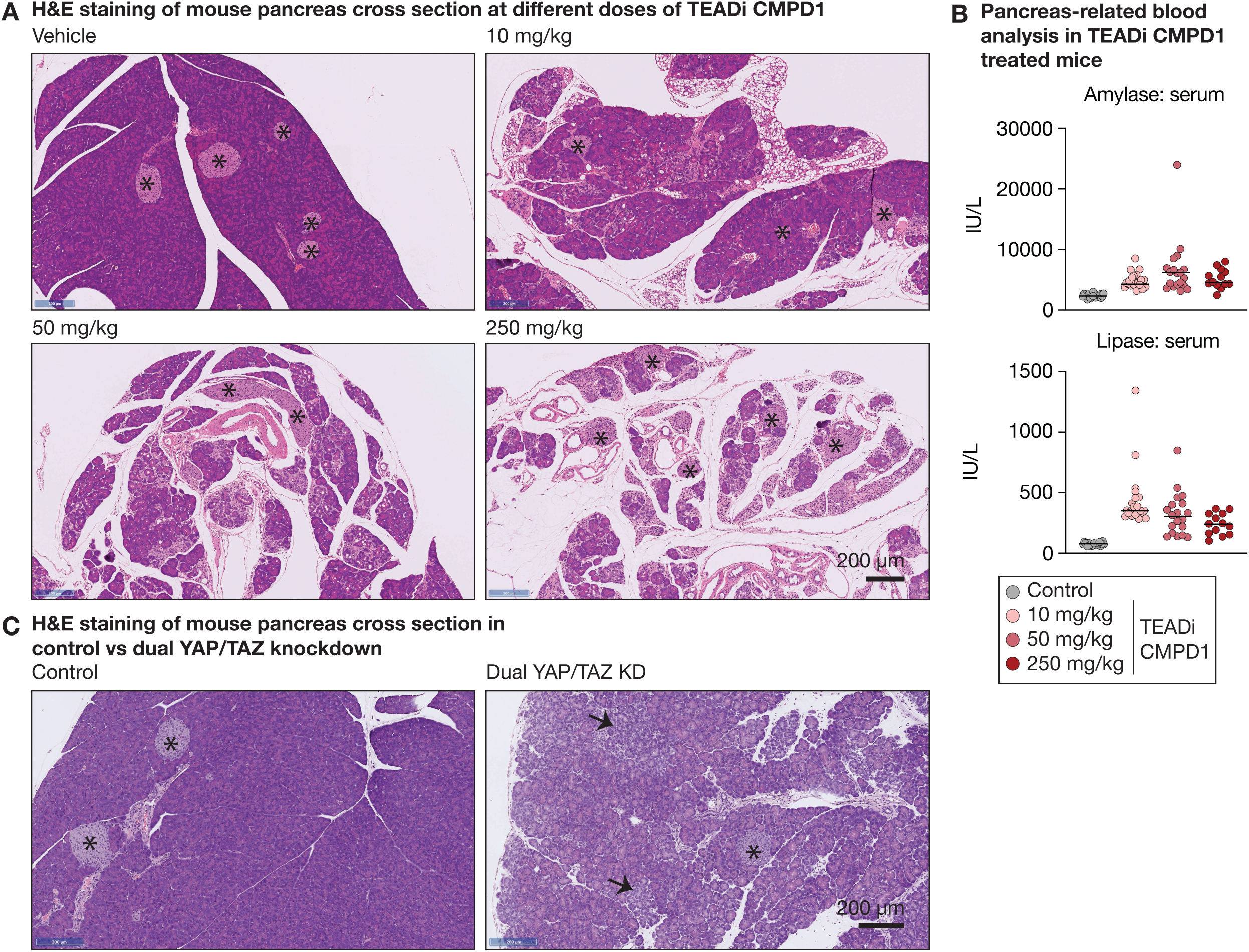
YAP/TAZ inhibition induces acinar cell atrophy in the pancreas. (A) H&E staining of pancreas cross-sections from mice showing degeneration and atrophy of exocrine pancreatic acinar cells after 28 days of vehicle or TEADi CMPD1 at 10 mg/kg and 50 mg/kg and after 22 days of TEADi CMPD1 at 250 mg/kg. Islets (asterisks) of the endocrine pancreas are unaffected. With increasing dose of TEADi CMPD1 there is increasing degeneration and eventual loss of pancreatic acinar cells. Scale bars: 200 µm. (B) Serum amylase and lipase concentration in mice after 28 days of vehicle or TEADi CMPD1 at 10 mg/kg and 50 mg/kg and after 17 to 24 days of TEADi CMPD1 at 250 mg/kg. Amylase and lipase are severely increased in all treated animals (≥ 10 mg/kg). As amylase and lipase are released from pancreatic acinar cells, there is a paradoxical relative decrease in the amylase and lipase concentration at 50 mg/kg and 250 mg/kg compared to 10 mg/kg, due to reduced functional pancreatic acinar mass. Dots represent individual animals; horizontal lines indicate the median. (C) H&E images of mouse pancreas from wildtype control and dual YAP/TAZ knockdown animals showing degeneration and atrophy of exocrine pancreatic acinar cells after 14 days of doxycycline administration. Islets (asterisks) of the endocrine pancreas are unaffected. In the dual YAP/TAZ KD mouse pancreas, acinar cells have paler staining and contain less zymogen granules, giving the pancreas a “moth-eaten” appearance. Scale bars: 200 µm.

### YAP/TAZ inhibition alters thymic architecture

To assess the impact of YAP/TAZ suppression on the thymus, we examined histological changes following pharmacologic or genetic inhibition. Histological examination revealed profound dose-dependent thymic lymphoid atrophy in mice treated with ≥ 50 mg/kg TEADi CMPD1 for up to 28 days and in transgenic mice with 14 days of doxycycline-induced dual knockdown of YAP and TAZ, with both models showing reduced cellularity of the thymus and loss of the normal microscopic distinction between the cortex and medulla (Figure 4A–B). Additional correlating findings with TEADi CMPD1 included reduction in thymus: brain weight ratio and circulating total lymphocyte counts (Figure 4C).

**Figure 4.**
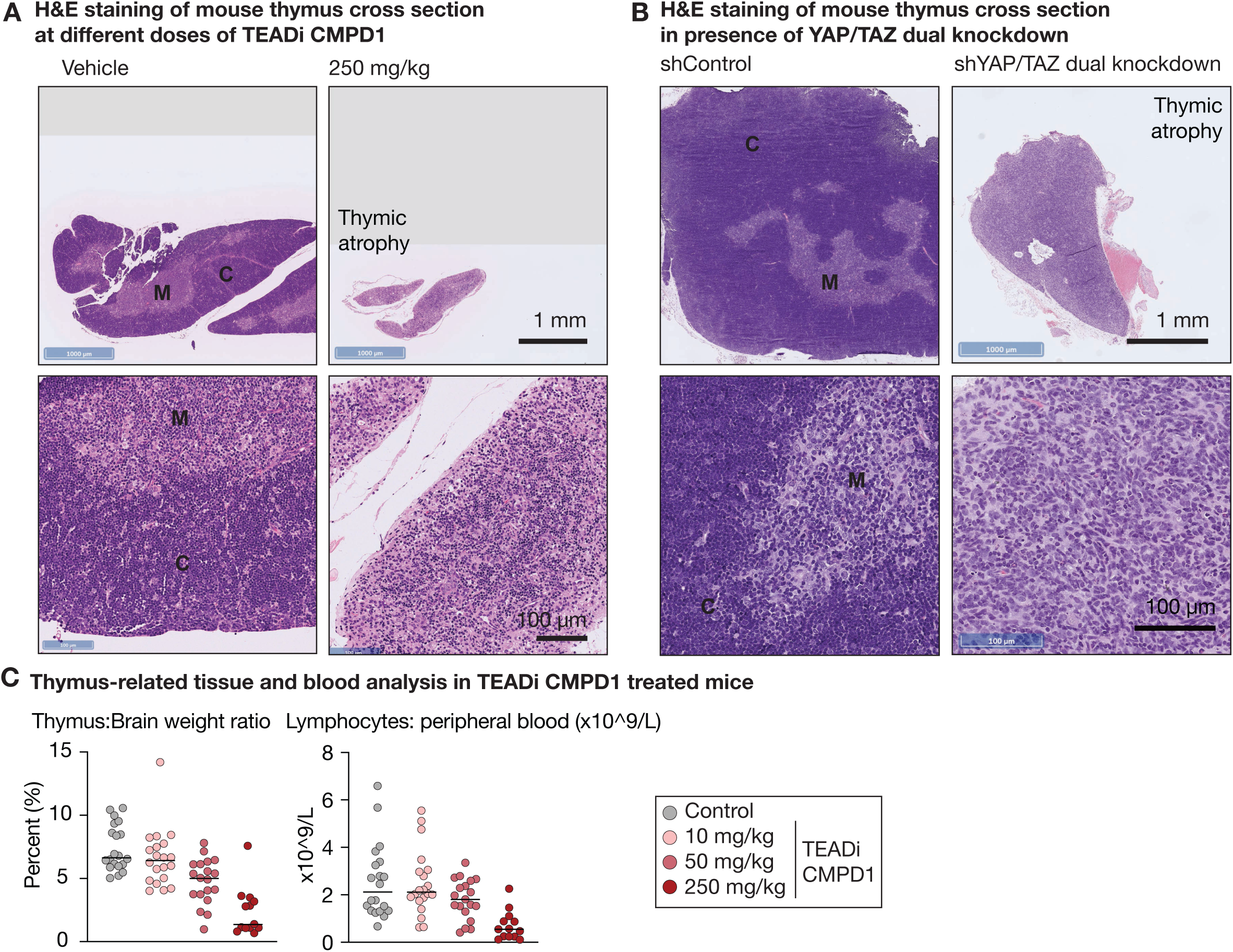
YAP/TAZ suppression causes thymic atrophy. (A) H&E staining of a transverse plane section of mouse thymus showing atrophy and loss of lymphocytes after 28 days of vehicle and after 22 days of TEADi CMPD1 at 250 mg/kg. In the 250 mg/kg dose group there is marked loss of lymphocytes from the cortex (C) and medulla (M) with loss of the normal corticomedullary demarcation. Scale bars: 1 mm and 100 µm. (B) Thymus weight and peripheral total lymphocytes count in mice after 28 days of vehicle or TEADi CMPD1 at 10 mg/kg and 50 mg/kg and after 17 to 24 days of TEADi CMPD1 at 250 mg/kg. There is a decrease in thymus weight (measured as the thymus to brain weight ratio to account for body weight loss) in the 250 mg/kg and, to a lesser extent, 50 mg/kg dose groups with a correlating decrease of circulating lymphocytes in the blood in the 250 mg/kg dose group. (C) H&E staining of a dorsal (coronal)-plane section of mouse thymus showing atrophy and loss of lymphocytes after 14 days of doxycycline administration to WT or Dual YAP/TAZ KD mice. In the Dual YAP/TAZ KD mice there is marked loss of lymphocytes from the cortex (C) and medulla (M) with loss of the normal corticomedullary demarcation.

### Reversibility of tissue damage following withdrawal of TEADi CMPD1

The renal and pancreatic findings noted at the end of the TEADi CMPD1 dosing period were irreversible and progressive during the 28-day recovery period. In the kidney, tubular dilatation in the corticomedullary region had progressed to tubular atrophy by the end of the recovery period and, along with a correlating decrease in kidney: brain weight ratio compared to both the end of the dosing period and the concurrent controls, indicated a permanent loss of renal tubules (Figure 5A). Urine volume and specific gravity, serum creatinine, and urea changes also did not fully recover, indicating a persistent reduction in renal function (Figure S2A). In the pancreas, there was increased severity of acinar cell atrophy compared to both the end of the dosing period and the concurrent controls, indicating progressive loss of pancreatic exocrine tissue. Islets were unaffected (Figure 5B). Although serum amylase and lipase elevations reversed, based on the histological appearance of the pancreas, this is more likely due to the atrophy-related, marked reduction in the pool of exocrine pancreatic cells from which amylase and lipase could be released, rather than any reversibility of the pancreas findings (Figure S2A). The alterations in thymic architecture and decrease in thymus:brain weight ratio noted at the end of the TEADi CMPD1 dosing period were fully reversible at 50 mg/kg and only partially reversible at 250 mg/kg during the 28-day recovery period (Figure S2A, B). The decrease in body weight recovered during the recovery period, with the dosed animal groups having a similar body weight to control animals by the end of the recovery period.

**Figure 5.**
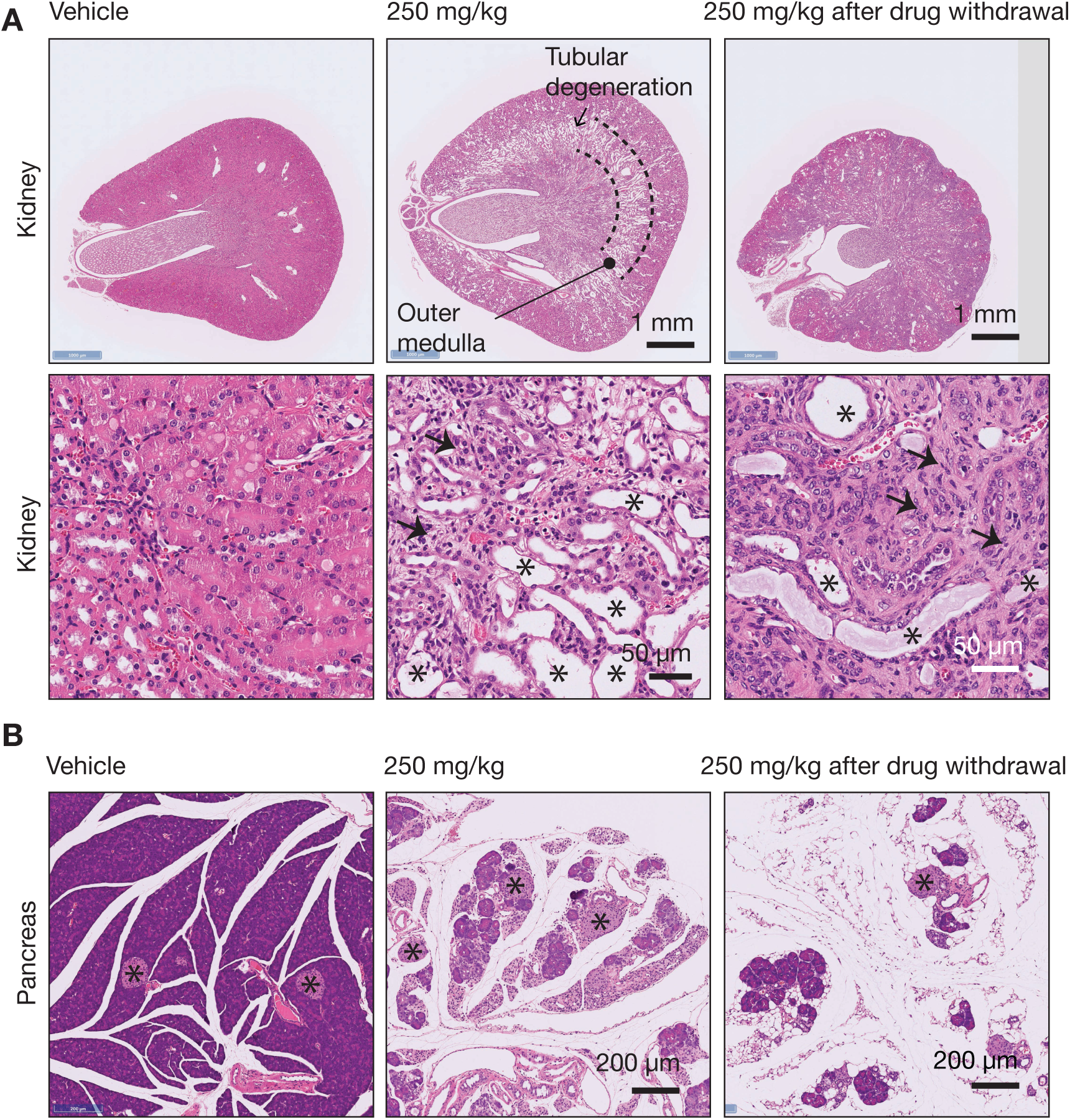
Irreversible tissue damage following withdrawal of TEADi CMPD1. A) (Top row) H&E staining of mouse kidney showing relative increase in size of the kidney, primarily due to dilatation and degeneration of tubules in the outer medulla, after 22 days TEADi CMPD1 at 250 mg/kg (center image) compared to vehicle (left) and a 28-day recovery (drug-free) period after 250 mg/kg TEADi CMPD1 treatment (right). The 250 mg/kg TEADi CMPD1 kidney after the 28-day recovery period (right) is smaller compared to both the vehicle kidney and the 250 mg/kg TEADi CMPD1 kidney after 22 days of treatment, indicating both lack of recovery and progression of kidney tubular damage and fibrosis during the drug-free period. Scale bars: 1 mm. (Bottom row) At the end of 22-days of 250 mg/kg TEADi CMPD1, there is tubular dilatation and degeneration and small amounts of early fibroplasia (arrow heads). After a 28-day recovery period, dilated and degenerate tubules (asterisks) are fewer in number and are replaced by increasing amounts of interstitial fibroplasia (arrow heads). Scale bars: 50 µm. The 250 mg/kg (22-day treatment) panels shown here are the same images presented in Figure 2A (kidney) and Figure 3A (pancreas) and are reused for direct comparison with the recovery cohort. B) H&E staining of mouse pancreas showing degeneration and atrophy of exocrine pancreatic acinar cells after 22 days of TEADi CMPD1 at 250 mg/kg (center image) compared to vehicle (left) and 250 mg/kg (right) TEADi CMPD1 after a 28-day recovery (drug-free) period. Islets (asterisks) of the endocrine pancreas are unaffected. The 250 mg/kg TEADi CMPD1 pancreas after the 28-day recovery period (right) has even greater reduction of exocrine acinar cells compared to both the vehicle pancreas and the 250 mg/kg TEADi CMPD1 pancreas after 22 days of treatment, indicating both lack of recovery and progression of exocrine pancreatic acinar cell atrophy during the drug-free period. Scale bars: 200um.

### Genetic suppression of YAP/TAZ perturbs chromatin accessibility and downregulates the aldosterone signaling Pathway in Kidney Epithelium

To investigate the molecular mechanisms of renal findings following dual YAP/TAZ knockdown (KD), target EPCAM+ kidney epithelial cells (Population II) were isolated using a sequential gating strategy to select single, live (7-AAD negative) cells. Within this population, GFP-positive cells were sorted as a reporter for doxycycline-induced shRNA expression, while GFP-negative cells served as wild-type controls. The efficiency of the knockdown was confirmed via mRNA quantification in the sorted populations. These isolated cells were then subjected to integrative ATAC-seq and RNA-seq analysis to correlate changes in chromatin accessibility with differential gene expression (Figure S3). ATAC-seq revealed widespread changes in the chromatin landscape, identifying a total of 15,210 regions with significantly reduced accessibility (closed peaks) and 1,594 regions with significantly increased accessibility (opened peaks) upon YAP/TAZ KD (Figure 6A and S4A, B; FDR<0.1, Log_2_FC>1). As expected, with loss of YAP/TAZ activity, motif enrichment analysis of the closed peaks (regions repressed in YAP/TAZ KD, Figure 6A) showed a significant enrichment for the TEAD transcription factor binding motif. This confirms that the observed loss of accessibility occurs at regions directly dependent on YAP/TAZ/TEAD transcriptional complexes. Other significantly enriched motifs included AP-1, FOX, P53, and ETS (Figure 6B). Conversely, the motif analysis of the opened peaks (regions induced in YAP/TAZ KD, Figure S4C) revealed enrichment for motifs associated with factors such as HNF (Homeobox), AP-1, NFκB, and nuclear receptor (NR) half sites (Figure S4D). To determine the functional relevance of the YAP/TAZ-dependent closing peaks, we annotated the differential ATAC peaks to nearest genes (+/-100kb around gene transcription start sites) and performed IPA (Ingenuity Pathway Analysis). This analysis revealed the genes and the enriched biological pathways that these repressed regions are proximal to, and might regulate, including the highly kidney relevant Aldosterone Signaling pathway and Rho family GTPase pathway which is known to be important for maintaining ion homeostasis and glomerular health (Figure 6C) (42) (43). This provided us with a potential link between YAP/TAZ activity and the observed renal tubular injury and proteinuria. Interestingly, IPA of genes linked to these induced regions showed enrichment for pathways related to Epithelial-Mesenchymal Transition (EMT) regulation (Figure S4E).

**Figure 6.**
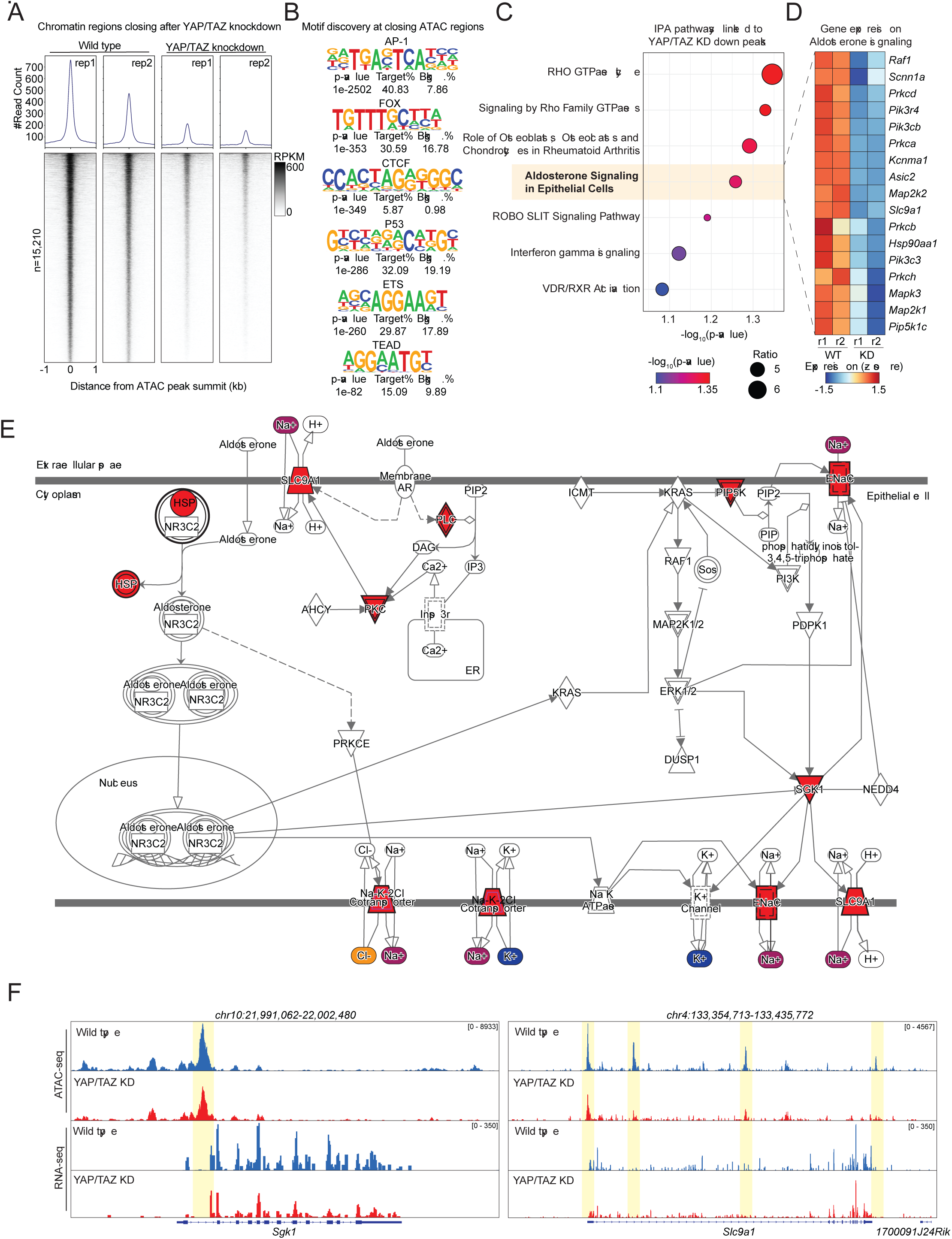
YAP/TAZ regulates the renal transcriptional program by directly controlling chromatin accessibility of genes involved in aldosterone signaling pathway. A) Density heatmaps and accessibility tracks (top) showing chromatin accessibility at 15,210 regions that significantly close (decrease accessibility) in kidney epithelial cells following dual YAP/TAZ knockdown (KD) compared to Wild Type (WT) controls. RPKM = Reads Per Kilobase Million. B) Top enriched DNA binding motifs identified by motif enrichment analysis within the 15,210 closing ATAC peaks. Enrichment is quantified by p-value. The TEAD motif is highly enriched, supporting a direct regulatory role for the YAP/TAZ-TEAD complex. C) Ingenuity Pathway Analysis (IPA) of genes annotated to the closed ATAC peaks. The plot shows the top canonical pathways that are significantly downregulated (negative z-score) or show enrichment (low - log_10_(p-value) on the x-axis), with the Aldosterone Signaling in Epithelial Cells pathway highlighted as the one of the most significantly enriched pathways that could provide insights into the kidney pathology observed in our mouse studies. D) Heatmap of RNA-seq expression levels (z-score) for a subset of genes belonging to the Aldosterone Signaling pathway. Transcriptional downregulation is observed across multiple key components upon YAP/TAZ KD (n=2 animals) compared to WT controls (n=2 animals). E) Schematic of the Aldosterone Signaling in Epithelial Cells pathway, highlighting core components that show reduced chromatin accessibility or transcriptional repression (red inverted triangles) following YAP/TAZ KD. Key effectors involved in sodium and water balance, such as Na^+^/H^+^ exchanger exchangers and Na^+^channel modulators, are affected. F) Genome browser tracks displaying ATAC-seq accessibility and RNA-seq expression (normalized counts, RPKM) at two critical loci: *Sgk1* and *Slc9a1*. Yellow shaded areas indicate specific ATAC peaks that close upon YAP/TAZ KD, correlating directly with decreased gene expression for both genes.

We further validated this pathway dysregulation using RNA-seq data from the same kidney epithelial cells. Consistent with the chromatin changes, the RNA-seq results confirmed transcriptional downregulation of genes involved in the Aldosterone Signaling pathway (Figure 6D). A closer look at the pathway schematic highlights that YAP/TAZ KD negatively impacts several key components which are critical for ion transport (Figure 6E). We highlight two major components of the Aldosterone-regulated sodium transport system that were consistently impacted at both the chromatin and transcriptional level. Visualization of ATAC-seq and RNA-seq data at the *Sgk1* (serum/glucocorticoid regulated kinase1) and *Slc9a1* (solute carrier family 9 member A1, a Na^+^/H^+^exchanger) loci confirmed that loss of chromatin accessibility at specific regulatory regions correlated directly with decreased gene expression upon YAP/TAZ KD (Figure 6F). Both *Sgk1,* a critical stabilizer of multiple epithelial ion pumps, and *Slc9a1,* a well-established Na^+^ transporter, are essential for regulating kidney epithelial ion balance, providing a potential molecular mechanism for the podocyte and tubular dysfunction observed in the YAP/TAZ-deficient mice (44, 45). Additional clinical pathology findings associated with the renal findings included increased serum sodium, chloride and potassium, increased serum creatinine and urea, decreased serum albumin with YAP/TAZ knockdown correlating well with reduced aldosterone signaling and proteinuria potentially due to impact on Rho family GTPase pathway (Figure S1D).

## Discussion

The Hippo pathway has gained significant recognition in oncology as a crucial regulator of tumor progression, prompting extensive efforts to harness its potential for therapeutic applications (2, 29). Given the important role of TEADs in normal physiology, concerns remain about targeting TEADs due to potential safety implications (32) (31). As several TEAD inhibitors are advancing through clinical trials, kidney-related adverse effects is emerging from both preclinical and clinical studies suggesting a possible “class effect” of pan-TEAD inhibitors (31–36). Recently, (37) reported cases of albuminuria and proteinuria in their VT3989 Phase I/II trial, representing the first proof-of-concept study targeting TEADs, in which these adverse events appear manageable, can be monitored via urine albumin/creatinine ratio, and are reversible with dose interruption or adjustment. While substantial progress has been made, ensuring patient safety remains a critical consideration throughout clinical development. In our study, we conducted a comprehensive pre-clinical profiling of both genetic and chemical perturbation of TEAD using transgenic mice and our in-house tool compound, TEADi CMPD1. We identified three key target organs and injuries that consistently emerged upon TEAD inhibition, regardless of the method used: kidney (tubular degeneration and proteinosis), pancreas (extensive acinar atrophy), and thymus (atrophy with lymphoid depletion). Pharmacologic TEAD inhibition produced greater adverse effects than genetic inhibition. A key difference could be the treatment duration, genetic inhibition was evaluated over 14 days, whereas TEAD CMPD1 dosing continued for 28 days, increasing cumulative exposure leading to severe phenotype. In addition, genetic perturbation may yield partial and/or tissue restricted target suppression, while systemic dosing likely drives broader and potentially deeper TEAD inhibition in sensitive tissues. Finally, compound specific liabilities (off-target activities) cannot be excluded.

Understanding the precise molecular mechanisms that drive adverse effects associated with pan-TEAD inhibition and the individual contributions of each TEAD paralog to this effect will be critical. Notably, TEAD1, TEAD3, and TEAD4 have been implicated in kidney function, particularly in maintaining podocyte and renal proximal tubular epithelial cell homeostasis (46–48). In addition, TEAD4 localizes to mitochondria, which are abundant in renal epithelium (46) and could also contribute to the renal findings associated with pan-TEAD inhibitors. As our understanding of the Hippo pathway deepens, additional roles for TEAD paralogs in kidney homeostasis may emerge, shedding light on previously unrecognized functions. Clinical trials with TEAD inhibitors, including VT3989 and IAG933 report proteinuria/albuminuria as common, albeit generally reversible, adverse events, (34–37), providing further support of a kidney-targeting class effect. So far kidney-related adverse events observed in clinical settings have been low grade and manageable through routine monitoring using standard renal biomarkers, such as serum creatinine and urine albumin levels, and adjustments to dosing regimens. Our findings extend this “class effect” by directly linking pharmacologic TEAD inhibition and genetic knockdown of YAP/TAZ to renal tubular degeneration, polyuria, and proteinuria. This phenotype is consistent with genetic models from previous studies demonstrating that Hippo pathway components are essential for renal homeostasis. Tubule-specific YAP knockout mice revealed increased urine volume and reduced urinary osmolality, attributable to impaired YAP-dependent expression of aquaporin, establishing the requirement of YAP for tubular function (49). Reginensi*, et al.* (50) show that conditional deletion of YAP in nephron progenitors in mouse kidneys leads to severely impaired nephrogenesis and abnormal nephron morphology. (51) reveal that in diabetic kidney disease, inactivation of YAP1 in renal tubule cells disrupts mitochondrial quality control, resulting in tubular injury, and elevated CXCL1-driven M1 macrophage infiltration. Podocyte-specific deletion of TAZ results in proteinuria, podocyte apoptosis, and progressive glomerulosclerosis (52). More recently (53), showed that YAP is also indispensable in podocytes, the combined loss of YAP and TAZ resulting in severe proteinuria and neonatal lethality. While glomerular abnormalities were not a prominent feature of the histopathology of kidneys in our studies, they may have developed in studies of longer duration had the tubular effects not been so severe. Based on our observations, it is possible that pathways affected by YAP/TAZ knockdown, including those involved in Rho GTPase regulation, may also contribute to the maintenance of nephron integrity. The degeneration of tubules primarily in the corticomedullary region of the kidney and correlating clinical observations of severely increased urine volume and electrolyte increases (sodium, potassium, chloride) with TEADi CMPD1 suggests a greater effect of TEAD modulation of the distal rather than proximal renal nephron in our studies which may be similar to the pathophysiological effect of impaired YAP-dependent expression of aquaporin in tubules as described by (49) and our demonstration of gene expression changes to the aldosterone signaling pathway in YAP/TAZ knockdown mice. Administration of TEADi CMPD1 for 28 days also resulted in progressive renal fibrosis, loss of renal tubules, and impaired renal tubular concentrating ability after a 28-day recovery period. Molecular analysis in our study revealed that YAP/TAZ KD in kidney epithelial cells leads to reduced chromatin accessibility and transcriptional downregulation of genes related to the aldosterone signaling pathway, which includes key components critical for ion transport. The enrichment for pathways related to Epithelial-Mesenchymal Transition (EMT) regulation may also have contributed to the development of the progressive renal fibrosis we noted in our TEADi CMPD1 studies.

In the pancreas we have seen striking exocrine pancreatic atrophy. The Hippo pathway is critical for maintaining pancreatic homeostasis as both hyperactivation and suppression of Hippo signaling in acinar cells destabilize pancreatic homeostasis (54). Inhibition of YAP has been shown to promote endocrine progenitor differentiation (55), while ectopic YAP induction in mature pancreas promotes ductal metaplasia at the expense of acinar cells (56) (57). During prenatal pancreas development, disruption of Hippo signaling upstream regulators like MST1/2 and LATS1/2 leads to a decrease in pancreatic mass and phenocopy of histological changes resembling pancreatitis (57, 58). However, little is known about the effects of YAP loss of function in the mature pancreas. Comprehensive reviews emphasize YAP and TAZ as central nodes coordinating proliferation, differentiation, and stress responses to maintain pancreatic homeostasis (54). More recently, (59) showed that acinar-specific deletion of Yap/Taz does not compromise basal pancreatic homeostasis, suggesting that YAP/TAZ are dispensable for acinar homeostasis in adulthood. However, our study shows a clear effect on pancreatic acinar cells via both genetic and chemical perturbation of TEAD with progressive atrophy of pancreatic acinar cells that do not recover in TEADi CMPD1 treated mice. A great deal remains to be understood about the systemic effects of YAP/TAZ loss of function, where it could render tissues highly vulnerable to injury, underscoring the need for a precise balance of YAP/TAZ activity to maintain acinar identity and resilience. These studies further highlight the importance of understanding the underlying mechanisms of action, and the physiological roles of the various members of the Hippo pathway.

In the thymus we have observed significant atrophy with systemic inhibition of YAP/TAZ. While scientifically interesting because of its role in T cell biology (60), its clinical translation and impact is to be determined, given the unclear role of thymus in humans adults as it naturally atrophies with age (61, 62). Thymic atrophy is also a very common stress response in mice that are suffering from adverse health effects and challenges in maintaining normal homeostasis as would have been occurring in our studies (63). Consequently, we are unable to determine if the effects of YAP/TAZ inhibition on the thymus in our study are due to TEAD inhibition, stress or, perhaps most likely, a combination of both.

After 28 days of treatment with TEADi CMPD1, findings of lesser severity were noted in some other organs, but it is unclear as to whether these findings are a result of TEAD inhibition alone or in combination with off-target inhibition of other targets and/or secondary effects related to stress and morbidity. Given the key role of the Hippo pathway in resistance to targeted therapy, this provides an opportunity to use the TEAD inhibitors as combination partners to targeted agents (15). These could involve longer term dosing in patients in combination with other pathway inhibitors, further highlighting the importance of a safe dosing regimen.

To address potential kidney related adverse effects in particular, many companies are shifting gears towards developing isoform-selective TEAD inhibitors which may potentially reduce kidney-related adverse effects (36, 64, 65). This approach could be particularly beneficial for patients with pre-existing conditions that compromise kidney function, ensuring safer and more effective treatment options. However, maintaining sufficient anti-cancer efficacy could be challenging in this context due to compensation by spared TEADs, and the development of irreversible toxicities will still be a concern, even in the context of advanced International Council of Harmonisation (ICH) S9 (cancer) indications (66). Another approach to improve the safety profile of TEAD inhibitors could be development of antibody-drug conjugates which preferentially target tumor antigens making disruption of homeostasis in normal tissues less likely. Growing interest in the Hippo pathway across oncology and immunology has expanded our understanding of its central role in development, tissue homeostasis, and disease. Insights emerging from ongoing genetic and pharmacological studies are beginning to define the toxicities a potential TEAD inhibitor should avoid while achieving meaningful clinical efficacy with an acceptable safety profile. With multiple small-molecule TEAD inhibitors advancing through preclinical and clinical pipelines, the field is entering an exciting phase focused on optimizing strategies to unlock the therapeutic potential of this critical signaling axis in cancer.

## Methods

### Mice

All mouse studies for transgenic or pharmacological modulation were approved by their respective Genentech, Inc or Shin Nippon Biomedical Laboratories, Ltd (SNBL) Institutional Animal Care and Use Committee. Animals were maintained under conditions of approximately 50% humidity (range 30% to 70%), appropriate room ventilation, and a 12h: 12h light: dark cycle, with free access to standard rodent food pellets and water.

Pharmacologic inhibition studies: Crl:CD1 mice were treated with TEADi CMPD1, a covalent palmitoylation-site binder, in a toxicity study for 28 days at doses of 0, 10, 50, or 250 mg/kg/day with a cohort of animals from the 0, 50 and 250 mg/kg groups followed for another 28 days after dosing cessation to assess recovery of any effects. TEADi CMPD1 was administered once daily via oral gavage.

Transgenic mouse studies: Conditional COL1A1.TRE.GFP.shYAPshTAZ; CAG.rtTA3 transgenic mice were used to achieve systemic knockdown of YAP and TAZ. In this model, TurboGFP (tGFP) together with shRNAs targeting YAP/TAZ were integrated ∼500 bp downstream of the Col1a1 3′ UTR by homologous recombination, under the control of a tetracycline response element (TRE), as previously described (41). To induce GFP expression and tandem YAP and TAZ knock down, mice were given 1 mg/ml doxycycline (Takara, 631311) dissolved in 5% sucrose water as the sole supply of water for 14 days after which the animals were euthanized and tissues collected.

### Ethics Statement

Our research complies with all the relevant ethical regulations. Animals were maintained in accordance with the Guide for the Care and Use of Laboratory Animals (National Research Council, 2011). Genentech and SNBL are both Association for Assessment and Accreditation of Laboratory Animal Care-accredited facilities and all animal activities in these research studies were conducted under protocols approved by the respective Institutional Animal Care and Use Committees.

### Immunohistochemistry

Knockdown efficiency was confirmed by immunohistochemistry using a custom antibody (clone SP519; Roche Tissue Diagnostics, Inc), which detects both YAP and TAZ. Antibody staining was performed at 0.1 μg/mL on the Ventana Discovery Ultra platform with standard Ventana reagents (Roche Diagnostics, Inc), including Ventana CC1 antigen retrieval, Ventana Rabbit OmniMap detection system, Ventana DAB chromogen and Ventana Hematoxylin II counterstain.

### Histopathology

Tissues were collected at necropsy and fixed in 10% (vol/vol) neutral-buffered formalin for approximately 24 h, and then routinely processed, embedded in paraffin, sectioned at 5 μm, and stained with hematoxylin and eosin (H&E). Stained sections were evaluated by a board-certified veterinary anatomic pathologist (M.L.).

### Clinical Pathology

#### SNBL

Blood was collected for hematology and clinical chemistry assessment via the caudal vena cava under isoflurane anesthesia prior to euthanasia for necropsy and analyzed for standard hematology and clinical chemistry parameters using the XN-1000V Hematology analyzer (Sysmex Corporation), BioMajesty Series JCA-BM6070 clinical chemistry analyzer (JOEL, Ltd), and Epalyzer 2 electrophoresis processing analyzer (Helena Laboratories Japan Co., Ltd.) Urine was collected after a period of 16 hours in a metabolic cage. Urine was assessed using Clinitek Advantus Urine Chemistry Analyzer (Kimball Electronics), a urinary refractometer (Atago Co. Ltd), and the BioMajesty Series JCA-BM6070 clinical chemistry analyzer (JOEL, Ltd).

#### GNE

Blood was obtained via cardiocentesis under isoflurane anesthesia prior to euthanasia for necropsy and analyzed for standard hematology and serum and urine clinical chemistry parameters using the XN-1000V Hematology analyzer (Sysmex Corporation) and AU480 automated clinical chemistry analyzer (Beckman Coulter Inc., Brea, CA), respectively. Urine was collected after overnight housing in metabolic cages.

### Kidney epithelial cell sort

Kidneys harvested from control and dual shYAP/TAZ knockdown animals were subjected to enzymatic dissociation to single cells. Then the kidney suspensions were washed in ice-cold PBS + 0.04% BSA, filtered (40 µm), and stained on ice in FACS buffer (PBS, 2% FBS, 1 mM EDTA) for 30 minutes protected from light with the following markers: GFP (native fluorescence; 488→525/50), CD45 (30-F11, BV605; 405→610/20), EpCAM/CD326 (G8.8, BV711; 405→710/50), and CD31/PECAM1 (MEC13.3, PE; 561→585/42; included in the panel but not used for downstream epithelial analyses); 7-AAD was added immediately before sorting per manufacturer’s recommendation. Antibodies were used in a 1:100 dilution. Cells were sorted on a FACS Aria sorter at low pressure with a 70 µm nozzle into cold PBS + 0.04% BSA using the following gate order: FSC/SSC, time gate, doublet exclusion (FSC-A/H; SSC-A/W), live cells (7-AAD⁻), then GFP⁺ and GFP⁻, then CD45⁻ EpCAM⁺ fractions from GFP+ and GFP-populations were collected separately for downstream ATAC-seq and RNA-seq.

### RNA isolation and library preparation

For each sample, 5–10 × 10³ sorted cells were pelleted (500 × g, 5 min, 4 °C), the supernatant was removed, and pellets were lysed in TRIzol (1ml). After 5 min at room temperature, chloroform (0.2 × TRIzol volume) was added, tubes were shaken for 15 seconds, incubated 2–3 min, and centrifuged (12,000 × g, 10 min, 4 °C). The aqueous phase was transferred to a fresh tube and RNA was precipitated with isopropanol (1 × volume) with Glycoblue (20 min, RT), pelleted (12,000 × g, 10 min, 4 °C), washed with 75% ethanol, briefly air-dried, and resuspended in 20 µL nuclease-free water. RNA quantity was measured with a Qubit high-sensitivity assay (Bioanalyzer trace optional at this input). Low-input cDNA and libraries were prepared using SMART-Seq V4 Ultra Low Input RNA kit following manufacturer’s protocol and sequenced as single-end reads (50bp).

### RNA-seq analysis

RNA-seq analysis was performed in R using HTSeqGenie (R package version 4.30.0) within the Bioconductor ecosystem to filter low-quality reads (≥70% bases with Phred <23) and remove rRNA/adapter matches (67). The remaining reads were aligned to the mouse reference genome (mm10) with GSNAP allowing up to two mismatches per 75-bp read (68). Gene-level counts were generated with featureCounts from the Rsubread suite using Ensembl gene models, and the resulting count matrix was carried forward (69–71). Genes with consistently low expression were removed with edgeR’s filterByExpr, libraries were normalized by TMM (calcNormFactors), and dispersions were estimated in a negative-binomial GLM framework (72–74). Differential expression was tested with the quasi-likelihood F-test (glmQLFit/glmQLFTest) and significance was defined at Benjamini–Hochberg FDR <0.05 with no fold-change cutoff (73–75). Gene annotations were obtained via biomaRt and mapped from Ensembl IDs to gene symbols for reporting and visualization (70, 76, 77).

### ATAC-seq

For each sample, 2,000-3,000 freshly sorted, viable cells were processed with the Active Motif ATAC-seq kit according to the manufacturer’s instructions. Nuclei were isolated, tagmentation was performed with Tn5 transposase for 30-minutes at 37C, and libraries were PCR-amplified (16 cycles) using kit indexing primers. Libraries were size-selected (SPRI) to enrich subnucleosomal fragments (∼100–200 bp), quality-checked by Tapestation, quantified by Qubit, and sequenced as paired-end reads (≥2 × 50 bp).

### ATAC-seq computational analysis

ATAC-seq computational analysis. Raw data were processed with the ENCODE ATAC-seq pipeline (v2.2.2) under default settings (78). Adapters were trimmed with cutadapt (v2.5) (79), reads were aligned to mm10 with Bowtie2 (v2.3.4.3) (80), and alignments were filtered for mapping quality and duplicates using samtools (v1.9) (81) and Picard (v2.20.7) (https://broadinstitute.github.io/picard/). Peaks were called with MACS2 (v2.2.4) using Tn5 offset correction and smoothing (82), and mm10 ENCODE blacklist regions were removed (83). Reproducibility was assessed with IDR (v2.0.4.2) (84), and standard QC included mapping statistics, library complexity, replicate concordance, and TSS enrichment. For visualization and summary profiling, bigWig coverage tracks were generated and normalized, and heatmaps and aggregate signal profiles for differential opening/closing peak sets were produced with deepTools (computeMatrix, plotHeatmap, plotProfile) (85). Differential accessibility was then assessed in R with DESeq2 modeling (86) in DiffBind (87) at FDR < 0.10; counts were generated on consensus peaks, depth-normalized, contrasts defined from a sample sheet, and results exported as induced (gain) or repressed (loss). Peaks were annotated with ChIPseeker using TxDb.Mmusculus.UCSC.mm10.knownGene and org.Mm.eg.db (88), and summaries were visualized in ggplot2. Motif enrichment on differential peak sets used HOMER with size-given, masked searches against a matched background (89).

### Homer de novo motif discovery

Motif enrichment was performed with HOMER (v4.11) on differential ATAC-seq peak sets (YAP/TAZ KD vs control), analyzing closing and opening peaks separately in mm10. For each test set we used the exact peak coordinates (-size given) with genome masking and compared against a background of length-matched consensus peaks not in the test set to control for GC and accessibility biases. HOMER reported both de novo and known motif enrichments; significance was assessed by Benjamini–Hochberg–adjusted q-values with a threshold of q < 0.05 (89).

## Data availability

Sequencing data generated in this study will be deposited in the NCBI Gene Expression Omnibus (GEO) and accession numbers will be provided upon publication. All other data supporting the findings of this study are available within the manuscript and supplementary materials, or from the corresponding author upon request.

## Acknowledgements

The authors would like to acknowledge members of the Genentech TEAD team, both past and present, particularly Ryan Ybarra, Melissa Schutten, Paula Katavolos, and Fiona Zhong. We also acknowledge the Genentech In Vivo Studies and In Vivo Pharmacology Groups, and the Genentech Core Research Pathology Facilities for their technical expertise and assistance with the execution of these studies.

## Author Contributions Statement

SP: manuscript writing and revision, figure preparation and data visualization, in vivo experimental protocol development and studies involving genetic perturbation of YAP/TAZ, single-cell isolation for transcriptomics study, interpretation of in vivo study results, data compilation and analysis of clinical pathology data.

ML: manuscript preparation and review, in vivo experimental protocol development, and analysis and interpretation of results from in vivo studies, including all pathology endpoints.

BD: manuscript review, editing, design of sorting strategy and epigenomic and transcriptomic data analysis. MS: performed ATAC-seq experiments.

JS, TH, SK, AD, RY, MN: manuscript preparation, editing, and review, in vivo experimental protocol development and experiment oversight, data compilation and analysis

---

**Figure S1.**
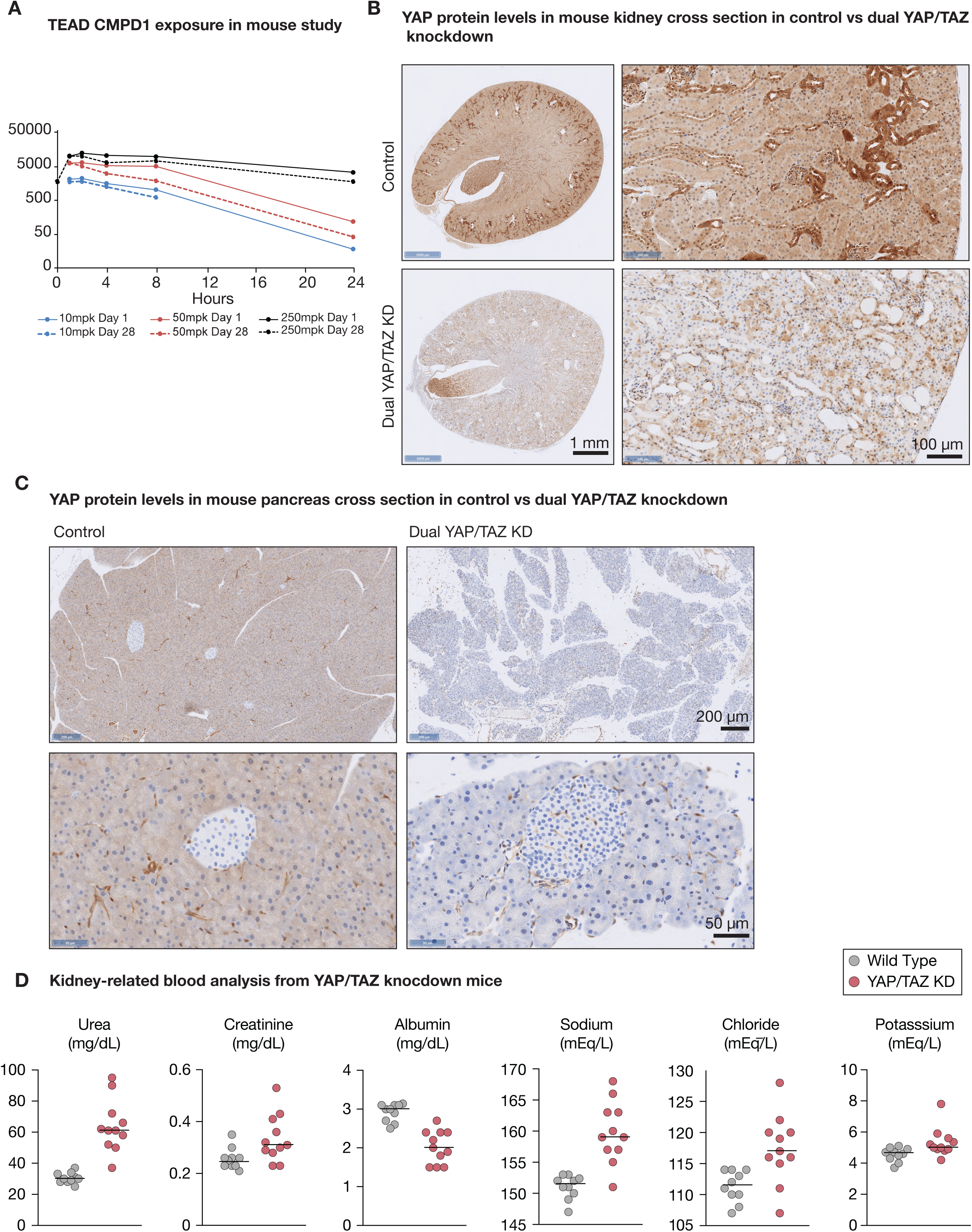
Characterization of dual YAP/TAZ knockdown mice. (A) Plasma concentration–time profiles of TEADi CMPD1 in mice dosed once daily for 28 days at 10, 50, or 250 mg/kg/day. CMPD1 exposure was assessed on Day 1 and Day 28, confirming dose-dependent systemic exposure. (B) Immunohistochemistry analysis of YAP/TAZ protein levels (brown staining) in kidney from control and doxycycline-treated mice with dual YAP/TAZ knockdown. Scale bars: 1 mm (top panels), 100 µm (bottom panels). (C) Immunohistochemistry analysis of YAP/TAZ protein levels (brown staining) in pancreas from control and doxycycline-treated mice with dual YAP/TAZ knockdown. Scale bars: 200 µm (top panels), 50 µm (bottom panels). (D) Kidney related blood analysis from wildtype control and dual YAP/TAZ knockdown mice after 14 days of doxycycline administration. Serum urea, and serum creatinine are increased, and serum albumin is decreased upon YAP/TAZ knockdown. Serum sodium, chloride and potassium are increased upon YAP/TAZ knockdown. Urine was not analyzed during this study. Dots represent individual animals; horizontal lines indicate the median.

**Figure S2.**
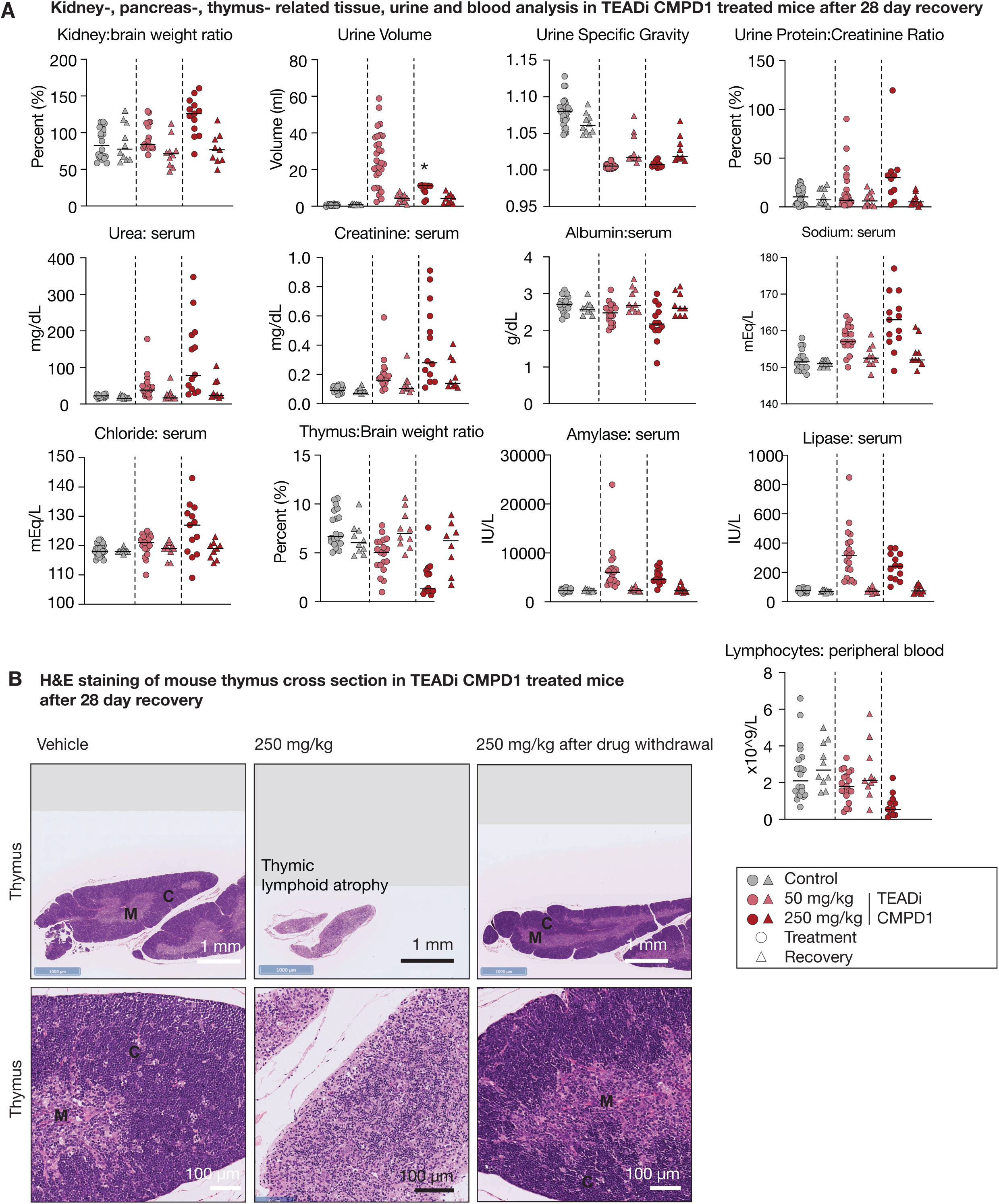
Kidney-, pancreas- and thymus-related tissue, urine, and blood analysis after TEADi CMPD1 withdrawal. (A) Kidney, pancreas, and thymus-related organ weights, urine and blood analysis in mice after 28 days of vehicle or TEADi CMPD1 at 50 mg/kg and after 17 to 24 days of TEADi CMPD1 at 250 mg/kg, followed by a 28-day recovery period showing only partial recovery to time-matched control levels for most parameters. As described in figure 2B, urine volume for dose 250 mg/kg (marked with an asterisk) could not be recorded correctly at the end of the dosing period because the container had overflowed. Dots represent individual animals; horizontal lines indicate the median. (B) H&E staining of a transverse plane of mouse thymus showing atrophy and loss of lymphocytes after 22 days of TEADi CMPD1 at 250 mg/kg (center image) compared to vehicle (left) and 250 mg/kg (right) TEADi CMPD1 after a 28-day recovery (drug-free) period. Thymic atrophy is partially improved after recovery with improving lymphoid cellularity and return of the demarcation between cortex (C) and medulla (M). Scale bars: 1 mm (top panels), 100 µm (bottom panels). The 250 mg/kg (22-day treatment) thymus panels shown here are the same image presented in Figure 4A and is reused for comparison with the 28-day recovery group.

**Figure S3:**
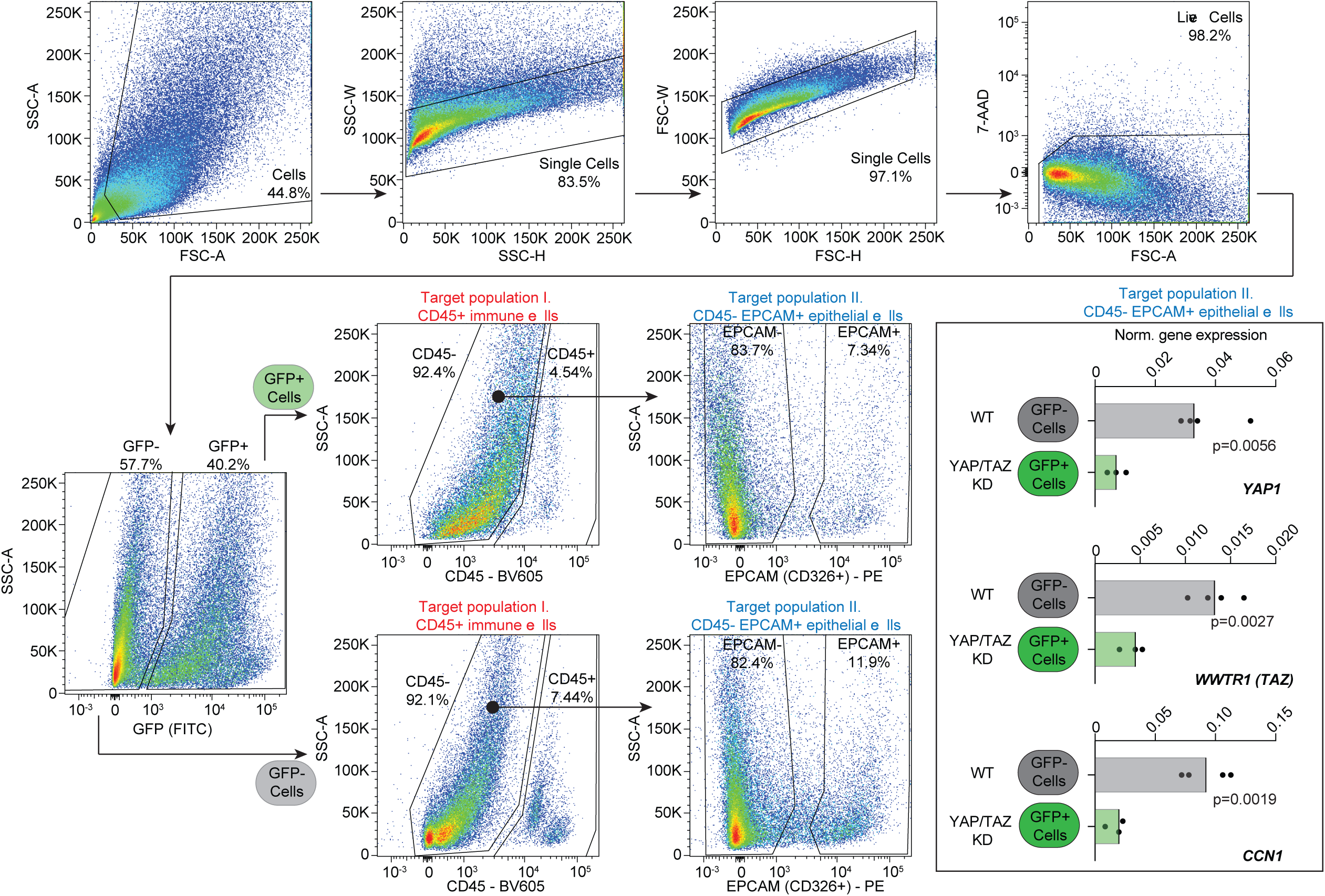
Sorting strategy for kidney epithelial cells. FACS gating strategy for the isolation of kidney epithelial cells from YAP/TAZ knockdown transgenic animals. Sequential gating was used to identify single, live cells (7-AAD negative). Sorting of GFP-negative (Wild Type) and GFP-positive (YAP/TAZ KD) cells within the target populations. GFP expression served as a reporter for shRNA induction following doxycycline administration. Target populations were isolated based on CD45-EPCAM (Epithelial cells, Population II) expression. Quantitative validation of knockdown efficiency in sorted (CD45-, EPCAM+) epithelial cells. Plots show normalized (to *Actb*) gene expression for *Yap1*, *Wwtr1* (TAZ), and the canonical Hippo target *Ccn1* (CYR61). Significant reduction of *Yap1* and *Wwtr1* expression is observed in epithelial cells with a corresponding decrease in *Ccn1* levels. p-values were calculated using an unpaired t-test. Dots represent individual animals.

**Figure S4.**
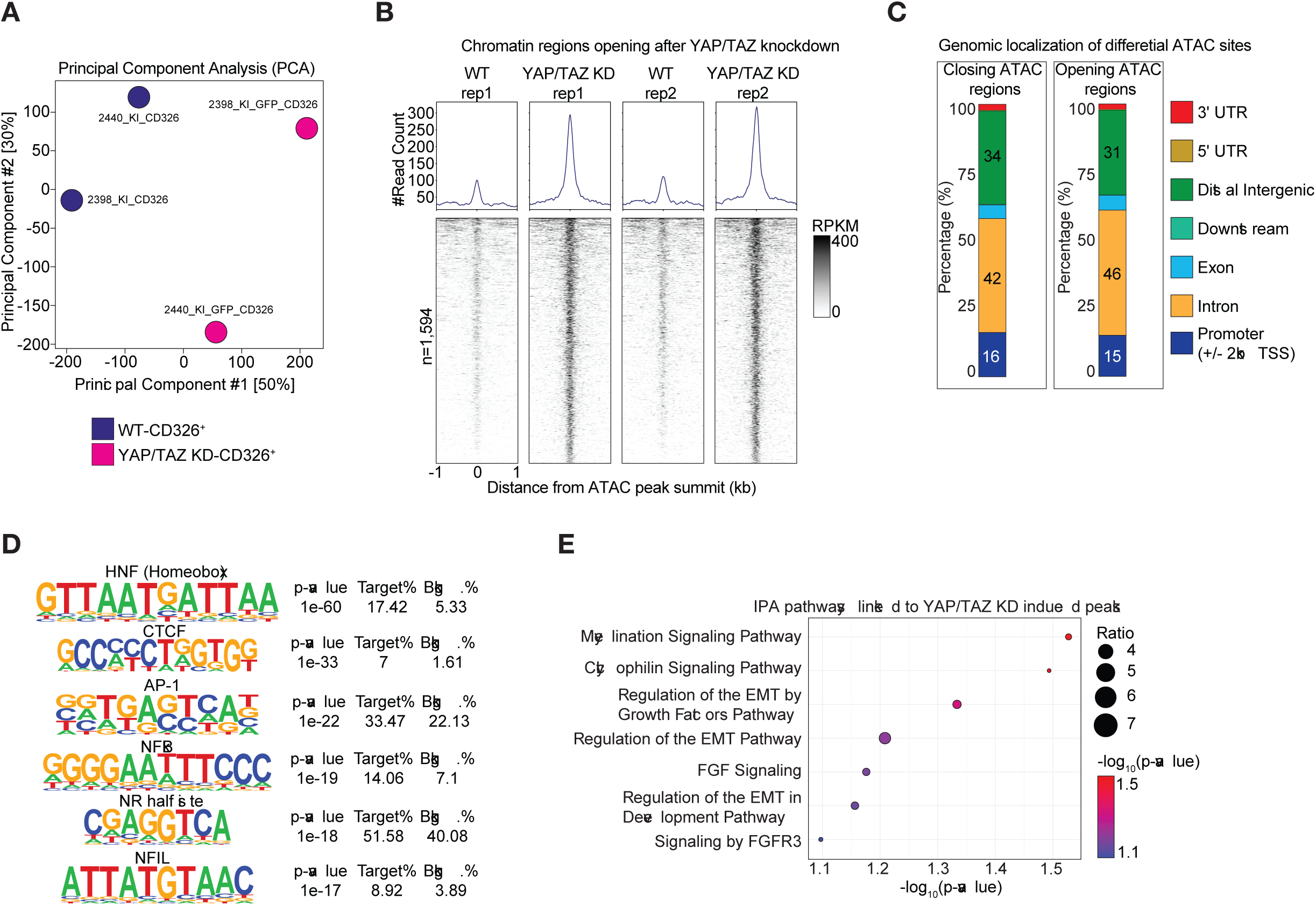
Analysis of Chromatin Regions Opening After YAP/TAZ Knockdown. A) Principal Component Analysis (PCA) of ATAC-seq data from kidney epithelial cells, showing clear separation of Wild Type (WT) and YAP/TAZ Knockdown (KD) samples, indicating significant overall changes in chromatin accessibility. B) Density heatmaps and accessibility tracks (top) showing chromatin accessibility at 1,594 regions that significantly open (increase accessibility) following dual YAP/TAZ knockdown compared to wild type controls. RPKM = Reads Per Kilobase Million. C) Genomic localization of differential ATAC sites (Closed and Opened regions) based on linear proximity to gene features (Promoter, Exon, Intron, etc.). D) Top enriched DNA binding motifs identified by motif enrichment analysis within the 1,594 opening ATAC peaks. Motifs include HNF (Homeobox), AP-1, NFκB, and NR half sites, suggesting the derepression of alternative transcriptional networks. E) Ingenuity Pathway Analysis (IPA) of genes annotated to the opening ATAC peaks. Pathways enriched include Regulation of the EMT Pathway and related signaling, suggesting that the loss of YAP/TAZ activity leads to the activation of gene expression programs associated with epithelial plasticity and mesenchymal characteristics.

